# Ion Channel Reaction Networks: Dielectric Screening and the Importance of Off-Pathway Flux

**DOI:** 10.1101/2025.03.06.637869

**Authors:** Hannah Weckel-Dahman, Ryan Carlsen, Alex Daum, Maxwell He, Tyler G. Southam, Jessica M.J. Swanson

**Author notes:** **Corresponding Author: Jessica M. J. Swanson** - Department of Chemistry, University of Utah, Salt Lake City, UT, 84112 – United States.

## Abstract

The transport of ions through channels involves multiple rare-event transitions through a web of interconnected intermediates. Extracting open channel mechanisms generally requires quantifying the relative flux through these intermediates in response to a range of electrochemical gradients. Although this is ideally suited to network-based representations like Markov state models (MSMs), the relative contributions from different pathways and the importance of network resolution remain open areas of research. Herein, we use a complementary approach called multiscale responsive kinetic modeling (MsRKM) to explore how the screening of ionic interactions and the competition between multiple mechanistic pathways contribute to channel mechanisms and current profiles of ion channels. We find that explicitly optimizing screened ionic interactions in the MsRKM framework vastly reduces the solution search space, enabling more efficient identification of physically robust solutions. Using a model of the Shaker Kv channel, we demonstrate that even when systems are well described by a single dominant flux pathway, the remaining contributing pathways and off-pathway flux play multiple essential roles, including shifting current profiles and mechanisms in response to different electrochemical gradients. We additionally discover that current continues to change above the experimentally predicted saturation point. Model systems explain how the degree of dielectric screening influences channel occupancy, the number of contributing pathways, and why current increases or decreases above its experimental saturation point. Our findings emphasize the importance of retaining a full network description to identify and understand ion channel mechanisms.

**Toc:** 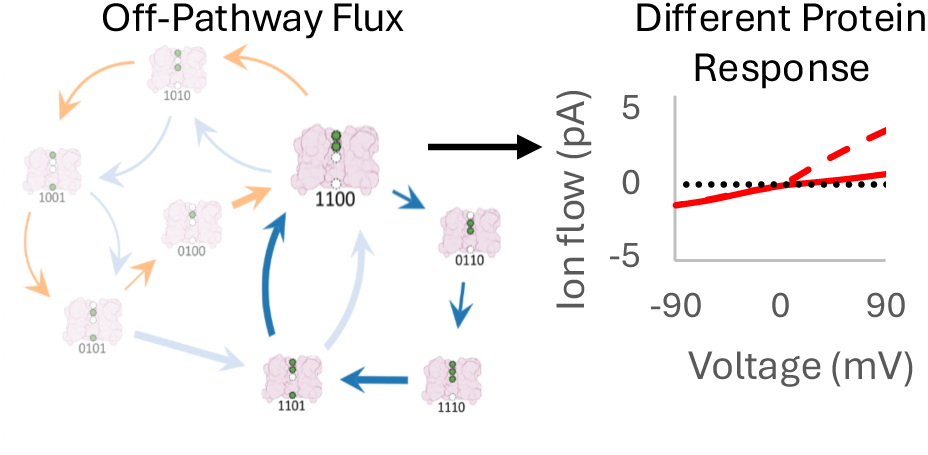

## 1. INTRODUCTION

Ion channels facilitate the selective passage of ions across cellular membranes, playing a crucial role in maintaining cellular homeostasis,^1, 2^ generating electrical signals,^3, 4^ and mediating various physiological processes.^2, 4^ These proteins have evolved to operate in highly dynamic environments with mechanisms that involve multiple rare events as ions transition between metastable intermediates (states). Markov state models (MSMs) have proven valuable in elucidating the complex state spaces involved in ion permeation, particularly in K^+^ channels.^5–8^ These models build upon the rich history of ion channel simulations,^9–11^ developing a network-based representation of nodes (states) and edges (transition probabilities) from simulations tracking ion permeation under applied electric fields. In principle, this network description provides a map of the full ensemble of protein-mediated ionic transitions. In practice, quantifying and characterizing the flux through a MSM network as a function of the applied voltage remains challenging. Recent work presented a mathematical framework to relate MSM transition probability matrices and associated flux matrices for short and long lag times to channel current.^12^ This corrects for issues of non-Markovianity in the former and the loss of less frequent path contributions in the latter. Although this required significant network simplification and was applicable to only one applied voltage, it produced the correct current directly from the MSM, which had not been possible previously. To complement this advance, a heroic effort combining MSMs with simulations run at multiple voltages demonstrated that tuning the electronic polarization, which led to increased channel occupancy, correctly reproduced complete IV curves for the first time, emphasizing the importance of properly balancing interactions in the force field.^8^

Given the complexity of most MSM network representations, a common question is whether the full network description is needed or if it can be simplified to a single or minimal number of contributing pathways. To date, ion channel MSMs have necessarily involved some degree of network simplification. The central challenge of network reduction is often identifying which nodes or clusters of nodes are the essential units of a network.^13^ Rapidly interchanging nodes may be combined or low-population nodes pruned, all with the goal of eliminating unimportant network dimensions while keeping particularly useful network properties consistent.^14–16^ In practice, this is a double-edged sword. While network reduction can enhance computational efficiency and clarify the dizzying array of data, it can also lead to significant deviations from experimental data^17–19^ when essential states and transitions are excluded.

Herein, we use a complementary approach, multiscale responsive kinetic modeling (MsRKM), to explore the roles of ionic repulsion versus a dielectric screening in the channel and the value of retaining a full network description, including both transport pathways with minor contributions to the net flux and off-pathway flux. MsRKM combines the strengths of bottom-up characterization from simulations with top-down refinement from experimental data to identify the continuous time transition rate matrix, as opposed to discrete time transition probability matrix, that best describes the process of interest. It was recently developed to describe electrochemically driven channels and transporters with rates that respond to chemical gradients and voltage.^17^ By including condition-responsive rates, MsRKM provides a network description of biomolecular mechanisms that can characterize the responsive flux under equilibrium, steady-state, and non-equilibrium conditions. Combining this with a reaction path analysis method based on Markov cycle theory that decomposes the flux into path contributions^20^ reveals how the reactive flux flows through this network under the electrochemical condition of interest. Most importantly, MsRKM provides a quantitative bridge between simulations (bottom-up rates) and experimental data (such as IV curves or flux assays), enabling data-based network refinement to identify kinetic solutions that are consistent with our best molecular level insight and macroscopic observations.

While MsRKM optimizations have identified critical network flaws and solutions that predict experimental data,^17^ reducing the kinetic solution space to physically robust and ultimately correct solutions remains the central challenge. Here, we describe an extension of MsRKM that explicitly includes screened ion-ion interactions in the optimization procedure. This drastically reduces the solution search space, more efficiently identifying a smaller number of physically reasonable solutions that can be grouped according to their dominant pathway. It also reveals the influence of ion-ion repulsions versus effective dielectric screening on channel occupancy and mechanism.

As with previous studies, a model of the Shaker Kv channel based on experimental data is used to probe the transport network. Solutions for this model are consistently dominated by a single pathway that contributes the majority of the flux (>96.5%), making it a seemingly ideal case for network reduction. However, we demonstrate that the remaining contributing pathways and off-pathway flux are essential in tuning the sensitivity of a protein to electrical versus chemical driving forces, in dictating if the relative pathway contributions shift under different electrochemical conditions, and in delineating how a channel responds to increasing ionic concentrations near its saturation point. This saturation behavior recapitulates previous experimental observations and theoretical work on the response of the ROMK1 potassium channel.^33,34^ We also reveal a surprising disconnect between experimentally determined saturation points for ion channels and complete network saturation with current that depends on the degree of dielectric screening. Collectively, these findings emphasize the importance of retaining a full network description. They also highlight the value of network properties in suggesting new experiments that could further refine the kinetic solution space, taking us closer to robust tools for reliable kinetic modeling.

## 2. THEORY AND METHODS

### 2.1 Implementation of Electrostatics in MsRKM

In top-down kinetic modeling, the optimization of kinetic parameters is likely to produce some solutions that violate physical principles that are not explicitly included in the optimization algorithm. Avoiding such errors is challenging in biomolecular processes with many potentially compensatory rates that may permit artificial flexibility in fitting experimental constraints. For example, in previous implementations of MsRKM on ion channels, solutions were often identified with networks that reproduced experimental data but with relative rates that were inconsistent with the expected ionic repulsions.^17, 21^ For example, solution 3 in Weckel-Dahman et al.^17^ fits the experimental data of the wild-type Shaker Kv channel, but it does so by predicting faster ion uptake rates with increased K^+^ occupancy (SI Table S2). It is important to note that it is possible for the impact of ion-ion interactions to violate the expected trends for relative rates. For example, the conformational changes induced by one ion binding could shield another binding site sufficiently to swap the relative rates. For highly dynamic systems, the range of potential impacts must be considered. For the remainder of the work herein, we develop kinetic models that fit the system before and after large conformational changes, such as the Shaker activation gate opening, and assume that the structure of the channel is consistent enough to ensure that rates speed up and slow down as expected due to like-charge repulsions.

In the absence of explicit constraints in the optimization procedure, the probability of obtaining a solution that consistently follows electrostatic trends is strikingly low. Given a linear channel that is selective to one ion substrate with N binding sites, the four binding/release rates are influenced by N-1 sites, while the 2(N-1) transfer rates are influenced by N-2 sites. This gives the following number of possible rate orderings:

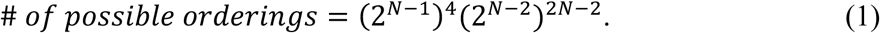

For the four-site channel studied herein, assuming rates are randomly sampled within their bounds, there is only a ∼1/16,777,216 chance of getting a solution with the correct rates. Thus, incorporating electrostatic constraints explicitly in the MsRKM optimization algorithm will drastically reduce solution phase space and eliminate a vast number of solutions that are physically inconsistent with our expectations for ion-ion interactions.

#### 2.1.1 Relating electrostatics to rate constants

Transition state theory provides a framework to relate the rate constant for a rare event transition to its activation free energy (ΔG^‡^):

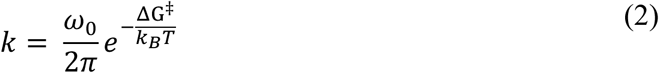

where *k_B_* is Boltzmann’s constant, T is temperature, ΔG^‡^ is the free energy barrier height, and *ω*_0_ is the attempt frequency. Although the attempt frequency will be transition-dependent, it has been consistently set to 1.16975×10^10^ ms^-1^ in this study to extract clear trends and insight.^22^ The influence of ion-ion electrostatics on the attempt frequency has been folded into the optimization of the effective dielectric factors, as described below. Future work will explore the impacts of electrostatic interactions specifically on the attempt frequency.

We now wish to compare rates for transitions involving only one ion in the channel to the equivalent transitions with more than one ion in the channel. Assuming that the transition free energy can be mapped onto a single reaction coordinate, such as the location of ion *i* along the z-axis (z_i_), the free energy profile of a single ion transition can be represented as:

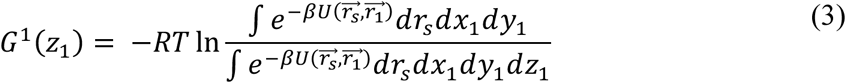

where R is the universal gas constant, β is the Boltzmann ratio, and U is the potential energy as a function of the cartesian coordinates of 3N degrees of freedom for a system of N atoms 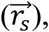 and ion 1 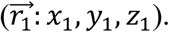 The addition of a second ion in the channel at a given binding site with position 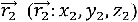 will shift the free energy profile due to the interaction of ion 2 with the system and with ion 1.^23–25^ This can be expressed as an additional term in the configuration integral including the Coloumbic interaction between the two ions:

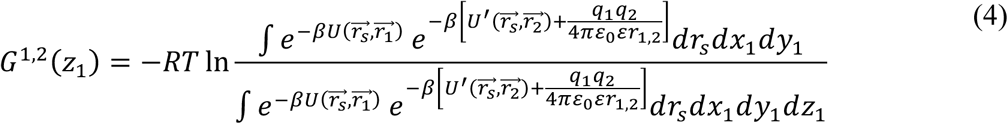

where *U’* is the potential energy of all atoms in the system 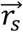 interacting with ion 2 at position 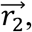 q_1_ and q_2_ are the charges of each ion, r_1,2_ is the distance between the ions, *ε*_0_ is the permittivity of free space, and ε represents the dielectric constant in the channel. Thus, the free energy profile for a two-ion system (Eq 4) will be shifted relative to the single-ion system (Eq 3). To properly capture this effect, the two free energy profiles must be fully sampled and directly compared.

Herein, we wish to limit our kinetic parameters to capture the correct trends in relative rates due to ion-ion interactions by approximating Eq 4 with 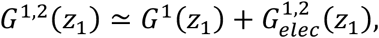 where 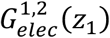 includes an effective dielectric screening parameter, *ε’*, that accounts not only for the screening in the channel environment but also for the shift in ensemble probabilities due to the interactions of ion 2 at a given binding position 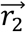 with the entire system:

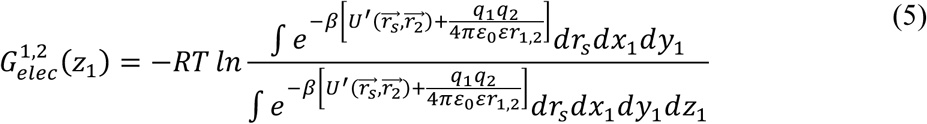

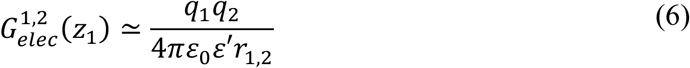

We then allow the optimization to determine the magnitude of *ε’*. As an optimized term, *ε’* is intended to include solvent screening, channel screening, and modest channel conformational changes induced by the occupancy of an additional ion. Major conformational changes, such as ion-induced gating that could counterintuitively stabilize proximal changes and would more substantially alter the underlying free energy surface, would not be effectively captured by *ε’*. A new kinetic model with distinct rates should be developed for protein responses of that magnitude.

Simplifying this expression, we see that the effective ΔG^‡^ for each transition is a summation of the activation free energy of the single ion occupancy and the shift in activation free energy due to the addition of ions represented as an effective electrostatic attractive/repulsive force:

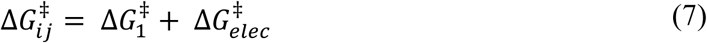

Similar to the activation free energy of a transition, 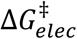 is the net difference in electric potential energy between the transition state and reactant states:

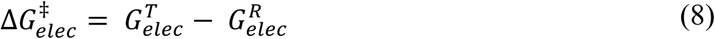

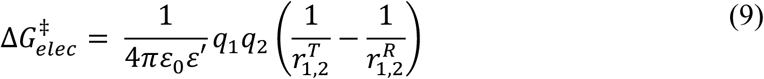

where 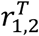 and 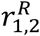 are the distance between an ion at the transition state and reactant position, respectively, and an ion at a different binding site within the protein. When the activation free energy is representative of a binding process, 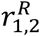 is taken at infinity to correctly represent bulk solution.

Combining equation 7 with equation 2, we obtain:

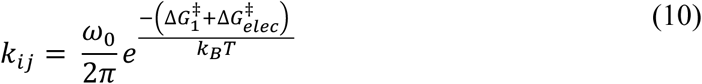

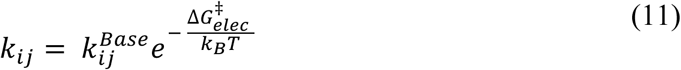

where 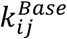 represents the transition rate from state i to j in a single-ion occupied channel.

#### 2.1.2 Ensuring correct electrostatic repulsion terms

During the optimization, the effective dielectric parameter *ε’* was allowed to range within reasonable bounds, from *ε*′=2 (permittivity of pure protein) to 100 (to allow for electrostatic screening higher than the permittivity of water). However, if each *ε’* was allowed to vary individually, relative trends would not be enforced. For example, ions occupying a site next to the transition could have a smaller impact on 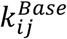 than ions occupying a site farther away. Although the simplest way to prevent this would be to define optimization search bounds for closer ions relative to those for farther ions, such a dynamic definition is not possible in Scipy. Thus, we instead use a product of scaling factors relative to the activation free energy for the farthest ion binding site.

For example, using the shorthand notation 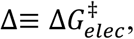 the activation free energy (Δ) for ion uptake from the intracellular side to site 1 in a four-site channel given additional ion occupancy at site 3 would be represented as:

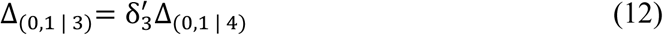

where Δ_(0,1 | 3)_ represents the activation free energy of binding at site one modified by ion occupancy at site three as the product of 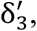 a scaling factor optimized for this transition, and Δ_(0,1 | 4)_, the activation free energy optimized for uptake to site 1 with only site 4 occupied. This ensures that the uptake to site 1 when site 3 is occupied is slower than or equal to that when site 4 is occupied. Similarly, 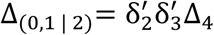 ensures that uptake with site 2 occupied is slower than or equal to that with site 3 occupied. In this sense, 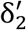 and 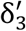 are multipliers relating the magnitude of influence of a closer ion to that of farther ions. If the transition moves an ion closer and repulsion should slow the transition down, the initial Δ_(0,1 | 4)_ would be positive, so *δ’* ≥ 1 would slow the transition down by making Δ_(0,1 | 4)_ more positive. A transition moving an ion farther away, such as Δ_(1,0 | 4)_, would be negative and *δ’* ≥ 1 will speed up the transition.

When more than one ion is present in the channel, impacts are cumulative (Figure 1). For example, if sites 2 and 3 are both occupied during uptake to site 1, then:

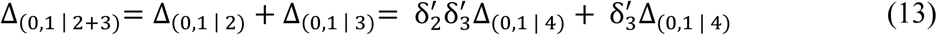

**Figure 1.**
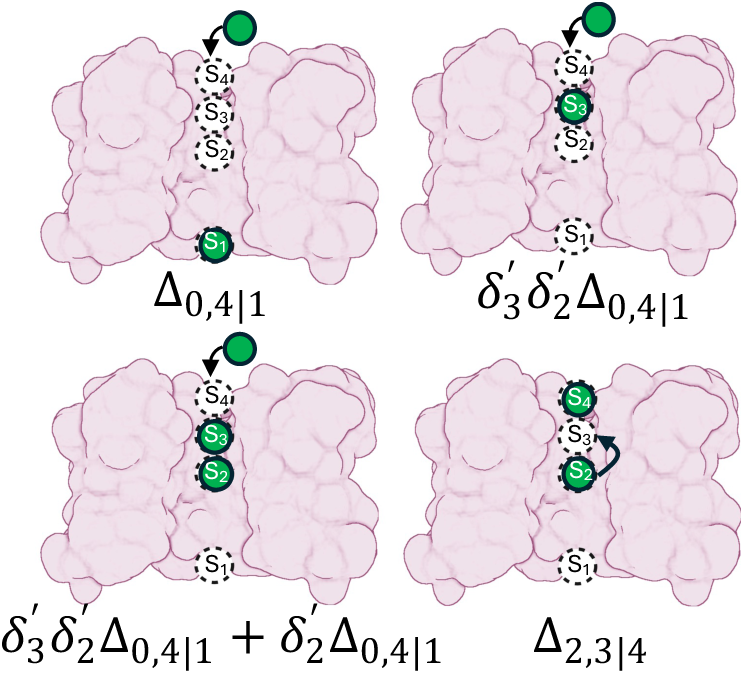
Delta multiplication descriptors for the specified transition given the specified occupancy.

As previously stated, the boundaries of each activation free energy were set to range between a lower bound for *ε*′ of 100 and an upper bound of 2 during the optimization process. The boundaries for each 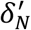 were set with the lower boundary for the multiplier being 1, reflecting protein conformational shifts canceling the closer electrostatic repulsion, while the upper boundary was set as the ratio between the free energies at a constant value of epsilon (SI Section 1.1).

### 2.2 MsRKM Model Construction and Optimization Protocol

#### 2.2.1 Model Construction

Three model systems were used in this work. Two of the models are based upon a 4-site model of the Shaker Kv channel used in previous MsRKM analyses^17^ (detailed in SI Section 1.1), one with electrostatics, herein called 4S-E, and the other without, 4S. The third system is a model 3-site channel previously used to extract trends in rectification^21^, with electrostatics implemented (3S-E). The intermediates of the 4-site Shaker models are defined by four K^+^ binding sites (S1, S2, S3, S4) as previously described^17^ (z^S1^ = -17Å, z^S2^ = 0Å, z^S3^ = 6.5Å, z^S4^ = 12.5Å) with all 16 states allowed (Figure 2 left) In the electrostatic model (4S-E), 10 transitions are base rates with only one ion in the channel. The remaining 46 rates were calculated by modifying the applicable base rate with the electrostatic deltas as described above. For the non-electrostatic model (4S), all 56 transition rates were modeled explicitly with no degeneracy assumed between the transitions. In the third model (3S-E), eight states were defined based on occupancy of the 3 cation-binding sites (Figure 2 right). The transitions include eight base rates and 16 delta modifiers.

**Figure 2.**
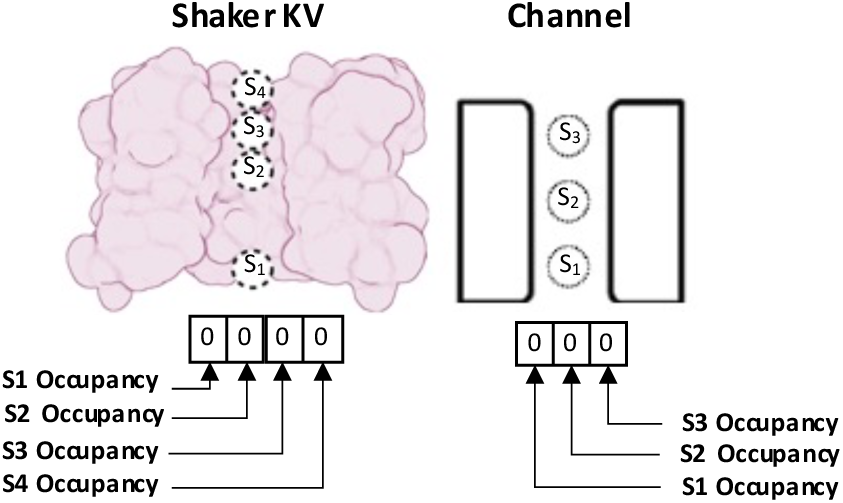
Binary state descriptors for the 4-site Shaker models (Left) and a generic 3-site channel (Right).

Both of the four site models utilized a symmetric coupling factor, consistent with previous studies^17^ (SI Figure S3 A), while the three-site model used a slightly modified symmetric coupling factor representing a linear voltage drop across the membrane^21^ (SI Figure S3 B). In the 4S and 4S-E models, the transition states were assumed to be at the midpoint between binding sites. In the 3S-E model, the transition state sites were initially located at the midpoint between binding sites or at the midpoint between an exterior binding site and the membrane boundary, but the locations were varied to explore rectification. For the site and transition state locations tested in the 3S-E model, see SI Table S1.

#### 2.2.2 Optimization protocol

The transfer rate coefficients of the four-site MsRKM models were given an upper boundary of 1×10^10^ ms^-1^ to represent the frequency of molecular vibrations, while the uptake rate coefficients were given an upper boundary of 1×10^8^ mM^-1^ms^-1^, which approximates the diffusion limit. All rate coefficients were given lower boundaries of 1×10^-10^ (mM^-1^)ms^-1^ to represent high-energy rare-event transitions.^26^ Delta boundaries were set as previously described. Boundaries were not determined for the 3S-E model as it did not undergo optimization.

The solution space of each model was explored with 100 randomly generated seed values. The Python library SciPy^27^ was selected to reduce the loss function using global and local optimization algorithms, with SciPy’s dual annealing as the global optimizer and Powell as the local optimizer. Each seed was used to initiate an optimization with 25,000 maximum global optimization searches. Each global optimization was evaluated by up to 100,000 steps of Powell, with all other hyperparameters set as recommended by SciPy documentation. The loss function compared the calculated flux of the MsRKM with experimental data for wild-type Shaker Kv channel^28^ while maintaining microscopic reversibility under equilibrium conditions (SI Section 1.1 for more details of the optimization protocol). Sets of rate coefficients satisfying our loss function cutoff were considered successful solutions if they deviated by less than 0.3 pA from experimental single-channel I-V curves. Solutions were considered unique if any individual rate constant differed by more than 2% from the corresponding rate constant of another solution.

### 2.3 Cycle Visualization

CycFlowDec^20^ was used for all models with a 500,000-step percolating algorithm with a burn-in of 4500 steps, a flux tolerance of 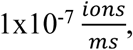 and an MRE cutoff of 1×10^-3^. Cycles identified with flux greater than 1×10^-3^ pA were selected for network visualization. Network graph visualizations of the cycles were prepared in Gephi 0.10.0 with the Polygon Shaped Nodes plugin installed. Nodes represent protein states, while edges represent the transitions between these states. Node sizes are proportional to the population of the respective states (See SI Equation S1). State populations are displayed next to each node where applicable. Edge weights and arrow sizes are proportional to the flux through the respective transitions (See SI Equation S2). Edge arrows indicate the direction of flux between states with different arrow colors representing different cycles within the network.

## 3. RESULTS AND DISCUSSION

### 3.1 Explicit inclusion of electrostatics reduces and refines solution space

A central challenge in kinetic modeling is narrowing solution space from the vast array of possible kinetic parameters that fit a limited amount of experimental and/or simulation data to those that correctly model the system. MsRKM was previously demonstrated to display predictive power for the Shaker K^+^ channel when refined with a sufficient amount of experimental data.^17^ However, none of the previously identified 500 solutions followed the expected trends of ion-ion electrostatic repulsion. In this study, the 4S wildtype (WT) Shaker model optimized with the same experimental data as previously used in addition to system saturation (see Methods) produced 98 out of 100 solutions from randomly generated seed rate coefficients (See SI Figure S1A). Despite each optimization including 25,000 global searches, the same solution was only found twice, resulting in 96 unique mechanisms. Once again, none of the solutions displayed the expected relative rates based on electrostatic interactions. This demonstrates the difficulty in obtaining physically reliable and thus physiologically relevant solutions through optimizations lacking explicit ion-ion repulsion.

When electrostatic interactions are included in the 4S-E model, 100 of 100 WT Shaker optimizations led to solutions. This time, equivalent solutions were identified 32 times, resulting in 77 unique minima. These 77 unique minima represent five groups based on their dominant pathways (See Figure 3), with the largest electrostatic repulsions causing decreased channel occupancy. For solutions with the same dominant pathways, we found that unique minima only differ in low flux or off-pathway cycles, which collectively contribute < 3.5% to the net flux. Interestingly, these mechanisms share a key similarity to previously identified mechanisms^17^: the dominant pathway comprises at least 96.5% of the net flux through the protein under the electrochemical conditions tested (-150 mV < Ψ < 150 mV, 43 mM < µ [K^+^] < 1.15 M). Whether single-pathway dominance is a universal feature of ion channels or specific to the Shaker Kv model remains to be determined. However, our three-site model results discussed below demonstrate that multi-pathway mechanisms are certainly possible. Collectively, the inclusion of electrostatic interactions in the optimization procedure appears to significantly limit the solution space at the mechanistic level while increasing the likelihood of finding a solution that reproduces experimental data.

**Figure 3.**
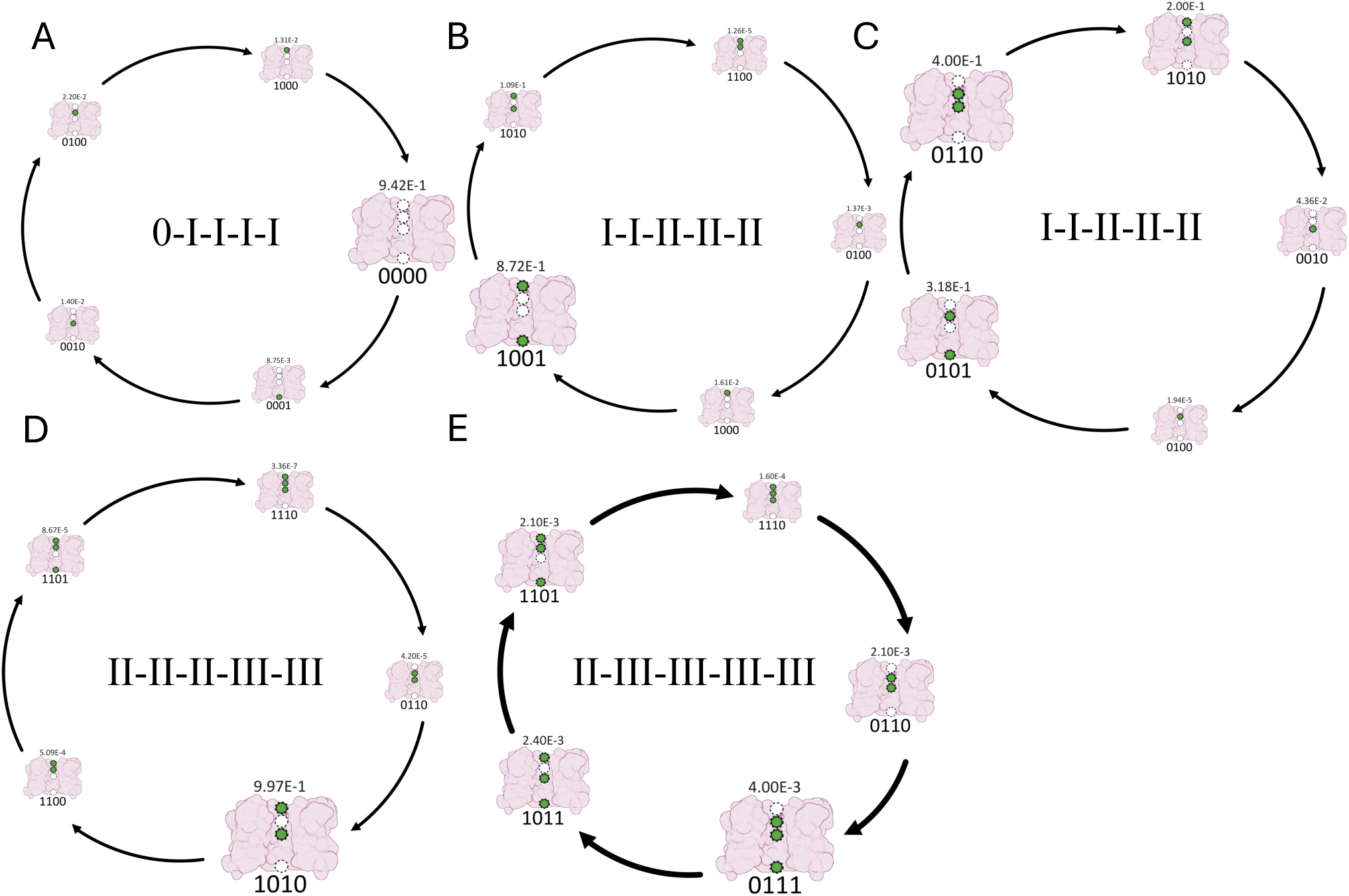
The 5 dominant cycles of the WT optimization. Roman numerals represent the ion occupancy of the states in the cycle. All cycles commence with the rightmost pictured intermediate.

For some systems, an additional metric that can be used to refine solution space is the location of the RLS. Previous work^17, 29, 30^ has suggested that the rate-limiting step (RLS) for low-conductance channels (like Shaker) should fall outside the selectivity filter. Of the 77 unique minima, only 24 had a RLS outside of the selectivity filter. Of these 24 solutions, the dominant pathways correspond to only 2 of the possible mechanisms shown in Figure 3. The most prevalent mechanism within these 24 solutions is similar to that proposed for KcsA simulations (Figure 3B)^31^, while the second most prevalent was identified in kinetic modeling (Figure 3E).^17^ If consistent with our previous findings, only these 24 minima can be possible Shaker solutions when reoptimized for the P457D Shaker mutant.^17^ However, the goal of this paper is not to define a solution that describes the Shaker Kv protein, but to understand the relevance of tracking flux through the full network of states; as such, all 77 solutions are analyzed to determine the roles of dominant and off-pathway cycles in modulating electrochemical response.

### 3.2 Off-pathway flux dictates unique electrochemical responses

Although each of the 77 solutions is dominated by one of the pathways shown in Figure 3 (≥96.5%), the non-dominant and off-pathway flux are still essential to capture the channel’s full electrochemical response. This is demonstrated by distinct network responses to electrochemical gradients not included in the optimization criteria. An extreme example of this is solutions 9 and 12, which both have the same dominant pathway (II-II-II-III-III, Figure 3D), the same flux limiting step (FLS) and rate limiting step (RLS) in that pathway, and the same secondary and tertiary pathways (SI Figure S4). Despite this they still react differently to changing chemical and electrical gradients (Figure 4A and B). Solution 12 is more responsive to a chemical gradient, while solution 9 increases flux more rapidly in response to voltage. They also have different responses to increasing symmetrical K^+^ concentrations driven at a constant voltage of 100 mV (Figure 4C). When flux is driven chemically solution 9 places more flux through the secondary pathway with increasing concentration gradients (SI Figure S4 Top), while solution 12 places more through the dominant pathway (SI Figure S4 Bottom). These two solutions are representative of a broader trend: even when solutions have the same primary, secondary, and even tertiary pathways, they still have unique responses to range of electrochemical gradients.

**Figure 4.**
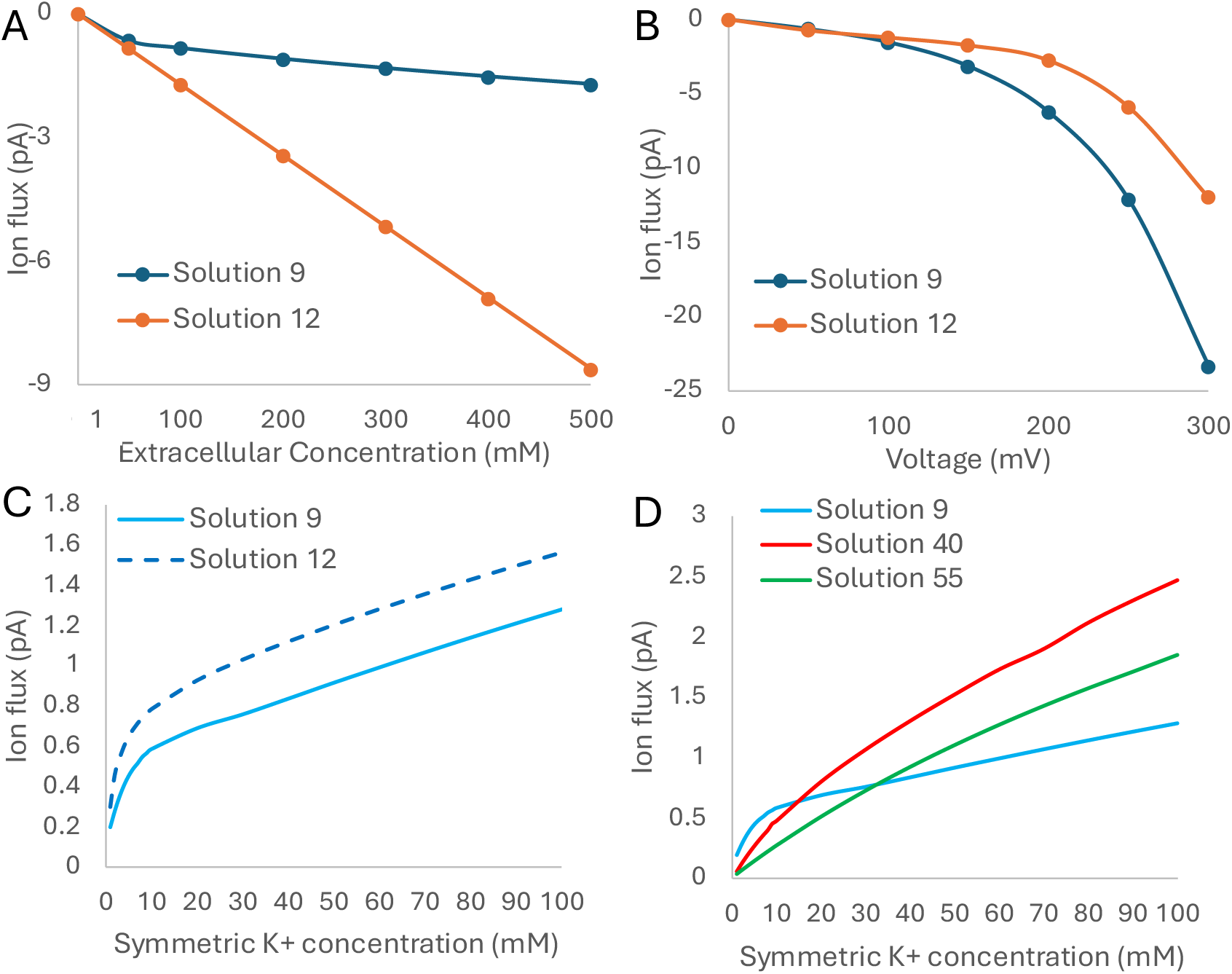
Comparison of two Shaker solutions with the same dominant pathway, highlighting the influence of off-pathway flux. Current was driven either (A) chemically by increasing [K^+^]_out_ while keeping [K^+^]_in_ at 1 mM, or (B) electrically by increasing voltage with symmetric 100 mM K^+^ concentrations. I-conc curves with increasing symmetrical K^+^ concentrations evaluated at +100 mV for (C) the same dominant pathway or (D) different dominant pathways.

Despite the above differences between two solutions with the same dominant pathways, mechanistic groups with the same maximum ion occupancies in the dominant pathway had similar I-conc curve shapes. Those with three or four ions in the channel (Figure 3 D, E), which corresponds to smaller electrostatic repulsions, show a more reserved increase in flux with increased K^+^ concentration (Figure 4 C, D blue). In contrast, mechanisms that allowed a maximum of one or two ion(s) in the channel (Figure 3 B, C, and A, respectively) show more pronounced responses to symmetric K^+^ concentrations (Figure 4D red and green, respectively). This suggests that solutions with larger electrostatic repulsions in their dominant pathway, which have lower channel occupancies, are more flexible in the the activation of new, higher-occupancy pathways, leading to greater overall flux at increasing concentrations. In contrast, solutions with smaller repulsions and higher occupancies in their dominant pathway have less flexibility to introduce new pathways and increase their net flux.

At the network level, we also see dominant pathway flux versus off pathway flux shift uniquely in response to increasing voltage. This is most notable when analyzing extreme voltage values where all mechanisms increase in flux but display different network behavior. For example in solution 9, the pathways shift congruently, maintaining the same relative contribution of all pathways to the net flux (Figure 5A). This differs from solution 55, which shifts its dominant mechanistic pathway at -500 mV (Figure 5B).

**Figure 5.**
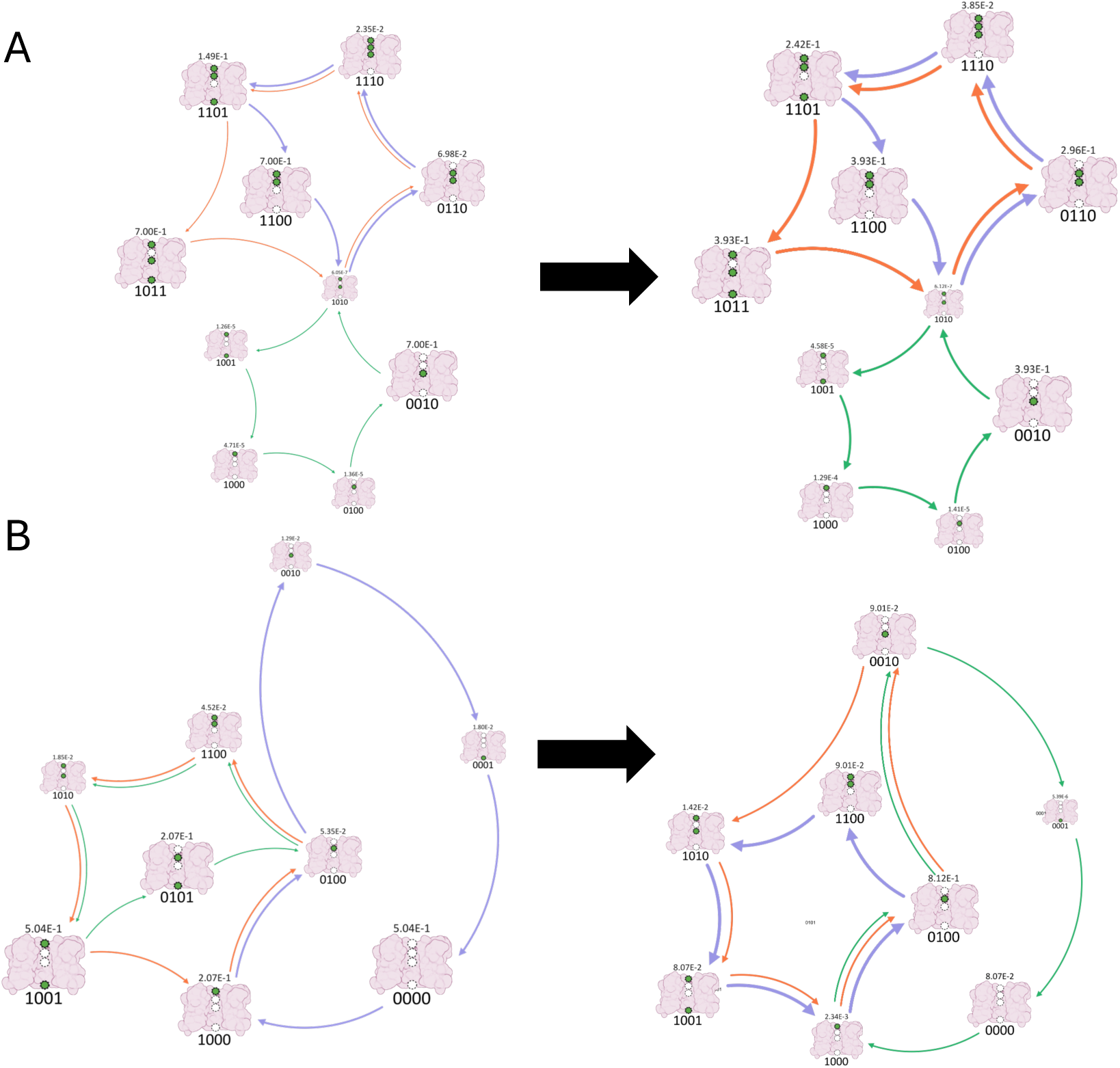
Mechanisms of the top cycles of solutions 9 (A) and 55 (B), evaluated at symmetrical 200 mM K^+^ concentration under -100 mV (left) and -500 mV (right).

Collectively, solutions that fit the same initial set of experimental data can have quite different network behavior in dominant, contributing, and off-pathway flux, endowing them with unique responses to electrochemical driving forces. This lends itself to both predicting new data and further refining solution space if correlative assays, such as those in Figure 3, can be done.

### 3.3 The role of off-pathway flux in experimentally observed saturation

The saturation point of a protein, often discussed in the context of enzyme kinetics, refers to the condition where increasing substrate concentration does not lead to an increased rate of product formation. For ion channels, this is the concentration at which net flux no longer increases. For the Shaker Kv channel, IV curves at 506 mM and 1.15 M are similar enough that the experimental saturation point (ESP) was determined to be 506 mM.^28^ From a network perspective, channel saturation occurs when the populations of all intermediates contributing to productive flux no longer change, leading to a constant flux despite increasing bulk ion concentrations. We call this the network saturation point (NSP). When analyzing the reported ESP in the 4S-E solutions, we found that only the dominant mechanistic pathway had saturated. As concentrations increased above 506 mM, the flux through previously noncontributing or minor pathways continued to change up to a much larger NSP. However, the changes in flux above the ESP are small. For example, the solution with the greatest discrepancy in flux, solution 3, only increases by 0.21 pA (8.86%) from the ESP at +/-90mV to the NSP of 7.1 M. Since this increase falls within typical experimental error^28^, the system would appear to saturate at the lower ESP concentration (506 mM).

Curious about the origin and significance of these network and flux changes above the ESP, we examined our 77 4S-E solutions for trends. Full network saturation varies between 7 M and 8.5 M (Figure 5A Green) with no clear correlation with the solution’s dominant pathway. To test how much the ESP and NSP could be altered, additional optimizations were run (SI section 1.1) allowing only non-dominant cycle rates to vary +/-10%. This revealed that networks with more flux coming from alternative pathways (defined by >2% of the net flux at the ESP) were capable of pushing their NSP higher (towards 9 M). Conversely, those with the smallest NSP (closest to the ESP) had the least contribution from alternative pathways (<1% of the net flux). However, the NSP remained greater than the ESP in all optimizations with the smallest devation being +80 mM. Tuning the non-dominant pathway rates also had a significant effect on the ESP— increasing it up to up to 2.5x (Figure S5A) or decreasing it to 400 mM (∼0.8x) (Figure S6B). Given that these impacts are far larger than the original non-dominant pathway contributions (<3.5%), this indicates that off-pathway flux can significantly shift the populations of dominant pathway intermediates where connecting rates were not allowed to change. Thus, the magnitudes of the ESP and NSP can both be significantly shaped by these non-dominant pathway network properties that are often removed or ignored during network reduction.

To unravel how the network properties were altered, we analyzed the mechanisms by which off-pathways could modulate the ESP. Two mechanisms were clear. In the first, the dominant path saturated at the same point as the original optimization, pushing all additional flux through minor and off pathways at the new ESP (Figure 6B). However, the dominant pathway retained dominance. In the second, minor and off-pathway contributions remained minimal, while the stability of the off-pathways containing intermediates of the dominant pathway increased. Consistent with our predictions based on the magnitude of changes possible for the ESP, this allows for altered flux through the dominant pathway through population modification (Figure 6C).

**Figure 6.**
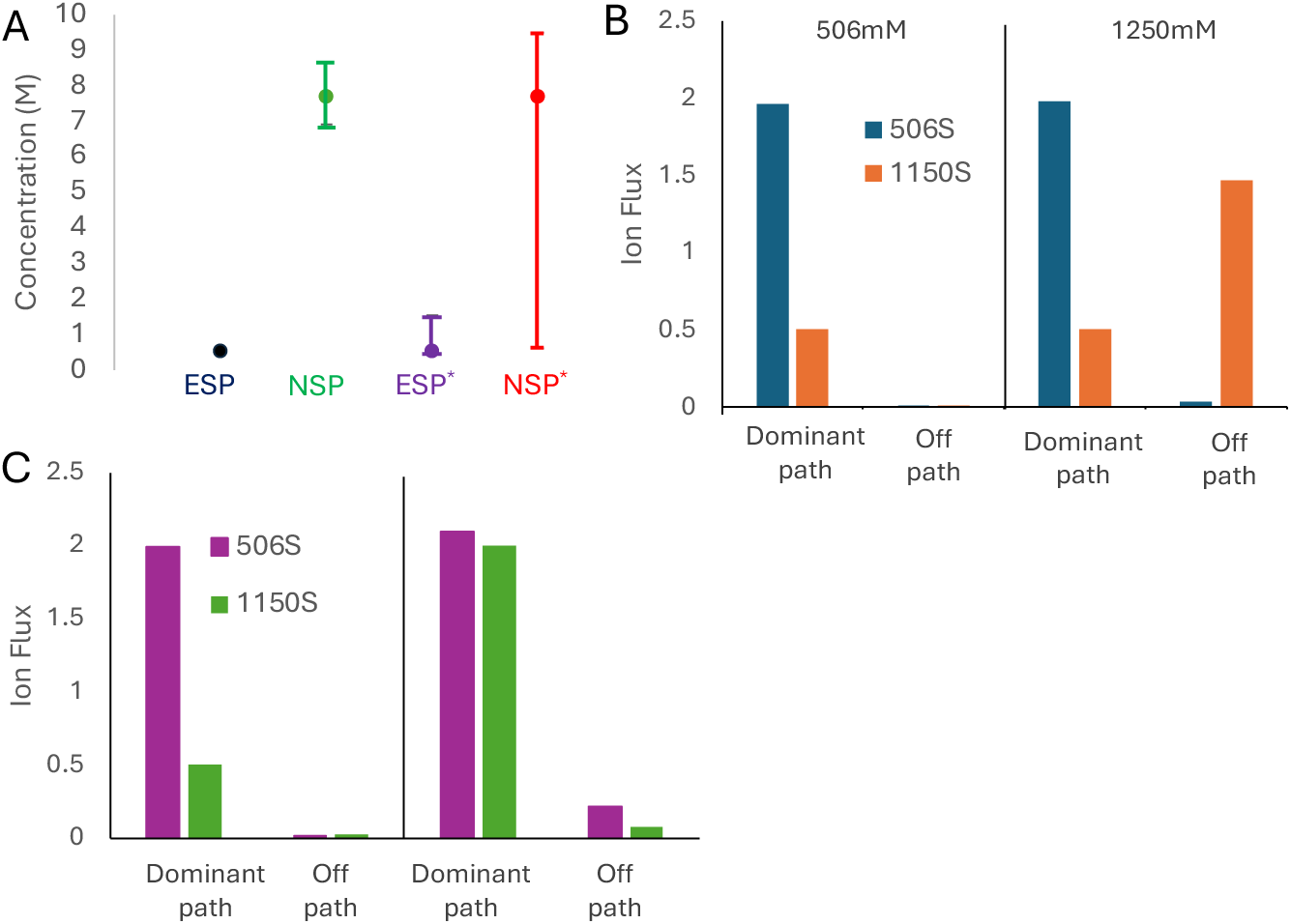
A) The range for the ESP (black) and NSP (green) of all 77 optimization solutions. The ESP* (purple) and NSP* (red) highlight the increased range in saturation points after varying off-pathway rates by ±10%. B, C) Comparison of the two mechanisms by which off-pathways contribute to the ESP. Solutions that saturate at 506 mM (blue & purple) were compared to those optimized to a ESP of 1150 mM (orange & green) evaluated at 506 mM (left) and 1250 mM (right), showing continued increasing flux can come from off-pathway contributions (B) or the dominant pathway (C).

Additionally, the magnitude of the flux at the NSP appeared to be correlated with the dominant pathway chosen. Solutions with one or two ions in their dominant pathway have a small increase in flux between the ESP and NSP (SI Figure S6 Dark Blue, Orange, and Green). However, solutions with three or more ions in their dominant pathways decreased in flux between the ESP and NSP (SI Figure S6 Blue and Purple). This was intriguing and dovetailed with further exploration using a model system described below.

Collectively, we find that the experimental and full network saturation points can be quite different and are influenced by the entire reaction network, even when over 96.5% of the flux comes from a single dominant pathway. Although the changes in flux between the ESP and NSP are small, assays focused on capturing these properties could provide a tremendous resource for resolving a system’s kinetic description. For example, 2D-IR has revealed the ability to sample the occupancy of ion channels,^32^ suggesting a potential route (comparing occupancy at different bulk concentrations) for the experimental detection of the NSP. This may allow experiments to either discriminate between model outputs or guide model optimization.

### 3.4 Introduction of electrostatics reveals a ‘sweet-spot’ for increased ion occupancy in channels

In order to further explore how the introduction of electrostatics influences saturation behavior, we used the 3S-E model to simulate the flux response to symmetric increases in concentration at different values of effective dielectric constant, ranging from vacuum permittivity (1) to an infinite magnitude to represent extremely large dielectric screening (Figure 7). Previously we had observed that increasing symmetric concentration above a given threshold eventually led to a decrease in flux,^21^ a consequence of the channel becoming fully occupied with ions and uptake outcompeting ion release. We replicate that trend (Figure 7) by setting our effective dielectric constant to infinity (representing perfect screening) but note that the trend still occurs for large values of the effective dielectric constant (ε=1000 representing strongly screened systems). Eventually, the increase in electrostatic repulsion resulting from a decrease in the effective dielectric constant (ε ≤ 100) prevents the channel from becoming completely crowded by ions, resulting in an asymptotic relationship between increasing concentration and flux.

**Figure 7.**
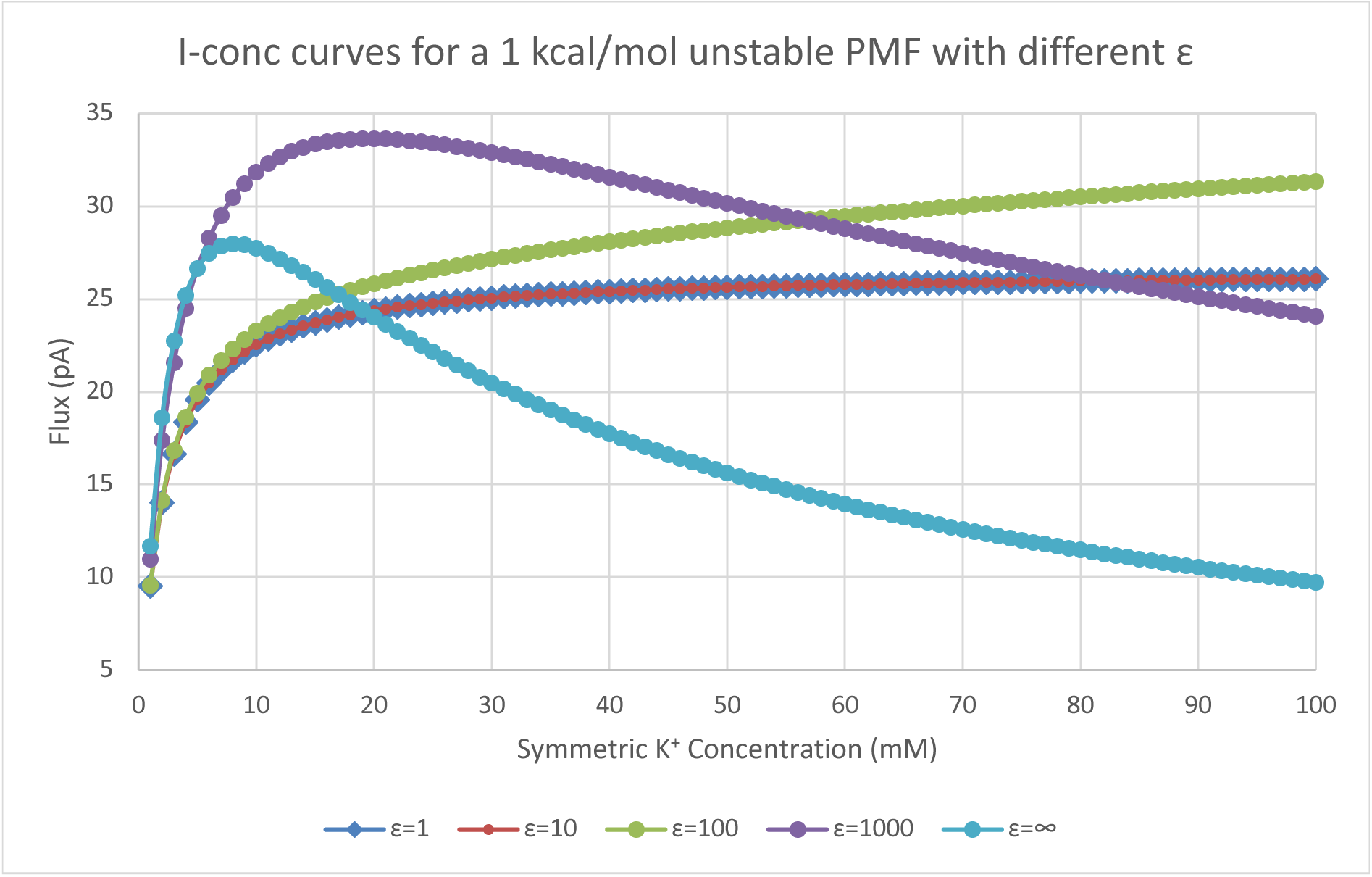
Flux through the 3S-E model with symmetric site locations and transition state locations as a function of increasing the intracellular and extracellular concentrations symmetrically. Binding site stability relative to bulk was set at 1 kcal/mol for 1 mM symmetric concentrations. Flux values were taken at +100 mV. Separate curves are flux values for different dielectric constants used to derive the electrostatic repulsion terms. Curves for dielectric constants ε = 1 and ε = 10 overlap.

Tying these observations about channel saturation and occupancy back to experiment, decreasing conductance above the saturation point has been observed and conceptually explained.^33, 34^ The most striking example is the ROMK1 channel, which exhibits a conductance maximum at only 300 mM and decreases its conductance by 75% at 1 M.^33^ The explanation for this behavior was elegantly captured in a simple 3-state model demonstrating that maximum conductance (*g*_max_) depends on transfer and release rates, while the concentration at which you hit *g*_max_ (X_max_) depends on the relative uptake rate. When uptake is fast relative to transfer and release, the channel will be increasingly blocked by uptake that opposes flux and thus conductance will decrease. The interesting addition demonstrated here is that channel tuning of the effective dielectric constant, modulating ion-ion repulsions, is essential. Since uptake to the final binding site when the adjacent site is occupied travels uphill against repulsions, while transfer from that adjacent site toward an edge site does not, larger electrostatic repulsions lead to increasing flux above the ESP.

We took the mechanisms of different values of the effective dielectric constant at three discrete concentration points (1 mM, 10 mM, and 100 mM) along the curve and analyzed them in detail. In these mechanisms, it becomes obvious that for low dielectric constant values (ε ≤ 10), the mechanism is reduced to a single pathway contributing to ion flux. This pathway is the stepwise motion of a single ion through the channel. While this pathway is also the dominant pathway at all dielectric constants for 1 mM symmetric concentrations, other significant pathways (responsible for > 5% of total flux) with higher ion occupancies begin to appear as the effective dielectric constant increases (Table 1).

**Table 1.** Number of significant pathways (>5% total flux) observed as a function of symmetric concentration (mM) and dielectric constant.

|  | Concentration (mM) |  |  |
| --- | --- | --- | --- |
| $\epsilon$ | 1 | 10 | 100 |
| 1 | 1 | 1 | 1 |
| 10 | 1 | 1 | 1 |
| 100 | 1 | 2 | 3 |
| 1000 | 3 | 6 | 3 |
| ∞ | 5 | 6 | 2 |

As expected, low dielectric constants favored dominant pathways with single-ion occupancy, while high dielectric constants favored dominant pathways with fully occupied channels. However, the secondary pathway of all mechanisms appears to be variable in ion occupancy, independent of the effective dielectric constant. For instance, at a dielectric constant of 100, there is a marked variation in the observed networks despite the same effective screening. These networks range from having a dominant pathway responsible for the majority of the flux, with less than 1% contributed by all remaining significant pathways, to networks where the dominant pathway contributes just over 50% of the flux. Interestingly, for each dielectric, the maximal flux corresponds with the largest number of contributing pathways. As concentrations continue to increase beyond this point, uptake increasingly outcompetes transfer and release—blocking the channel and eliminating paths with low ion occupancy, thereby decreasing the number of contributing paths. These results demonstrate that electrostatic interactions play a crucial role in controlling ion channel saturation and flux behavior by shifting the balance between transfer, release and uptake, and concomitantly the dominance of pathways with increased or decreased ion occupancy.

### 3.5 Introducing electrostatics minimally alters rectification trends

We previously explored the origins of rectification using a three-site model (3S) without electrostatics.^21^ As we introduced electrostatic constraints into our model, we revisited the same series of tests with 3S-E using a dielectric constant of 20. For all shifts in site and barrier location, electrostatic results are congruent with previous results, with only minor changes in magnitude. Similar rectification results were also observed for altering the barrier height, although there were more significant differences in the magnitude of these effects. However, the trends for changing the depth of the first binding site changed. In both low and high concentration cases, rectification was abolished when electrostatic repulsion was introduced.

This aligns with our initial explanation of the trends for rectification in the non-electrostatic model. Here, our rectification argument depended on additional ion occupancy in the channel setting up a competition between transfer and uptake. With low concentrations, when other ions cannot occupy the channel due to electrostatic repulsion, the competition between binding and transfer at the first site is abolished. With high concentrations, rectification is similarly abolished by this lack of competition. It should be noted that this lack of rectification is expected to be highly dependent on the concentrations and on the electrostatic screening. However, in our limited recapitulation of our previously reported rectification patterns, we find it significant that all are retained when electrostatic repulsions are included except those that depend exclusively on multiple ion occupancy.

## CONCLUSIONS

We have presented an extension of the MsRKM framework that explicitly includes ion-ion interactions in the optimization algorithm. This reveals the balance between like-charge repulsions and dielectric screening by the channel environment. It additionally eliminates a vast number of solutions that lose the proper relative rate relationships from the parameter search space. We find that the inclusion of electrostatics also limits the solution space at the mechanistic level while increasing the likelihood of finding a solution that reproduces experimental data. Thus, it allows for faster identification of physically reasonable solutions. For the model of the Shaker Kv channel studied herein, solutions could be grouped into five dominant mechanistic pathways. These dominant pathways reflect the effective electrostatic screening of the network as a whole, with higher channel occupancy (containing 3-4 ions) correlating with greater dielectric screening and similar I-conc curve shapes (Figure 3D). Interestingly, the non-dominant pathways of each solution appear to modulate how responsive the net flux is to chemical gradients and voltage (Figure 3A and B), even in solutions that share the same dominant pathway (Figure 3C). These non-dominant pathways also appear to influence electrochemical response of a protein at the mechanistic level.

Previous studies using MSM of ion channels under applied voltages have revealed voltage-induced mechanistic changes.^8,35^ While our results support this for some solutions (Figure 5B), we also reveal solutions that remain unchanged under increasing voltages (Figure 5A). In these solutions, the pathways of the network shift congruently, maintaining the same relative contribution of all pathways to the net flux. Suprisingly, even solutions with virtually identical dominant pathways and similar contributions to the net flux from non-dominant pathways but with different supporting networks, can display different mechanistic changes not only at different voltages but also at increased chemical gradients (SI Figure S4). Together, the relationship between flux, dominant pathways, and electrochemical conditions demonstrates that the entire network has a role to play in channel behavior.

We also find that the remainder of the network outside of the dominant pathway plays an important role in determing the saturation point—both for the entire network (NSP) and for that measured experimentally (ESP). Not a single solution is identified that saturates at the ESP. Rather, flux through the dominant pathway plateaus while other contributing pathways continue to change the flux up to the NSP. However, the magnitude of the current change is small enough that it would be difficult to detect without experimental effort directly focused on this behavior.

Whether the net flux increases or decreases above the ESP appears to be a function of the dominant pathway. Mechanisms with increased ion occupancy in their dominant pathway (high dielectric screening) show a decrease in the flux between the ESP and NSP, while mechanisms with two or fewer ions (stonger ion-ion repulsions) gradually increase flux between the ESP and NSP (SI Figure S7). This can be reconciled by considering the competition between forward movement of an ion from an edge-adjacent site into an edge binding site versus reverse flux uptake from solution into that site. Consistent with experimental observations and previous explanations, when uptake outcompetes transfer, conductance above a saturation point will decrease.^33,34^ What was not previously discussed is that ion-ion repulsions play a direct role in this balance. When screening is high, uptake can outcompete transfer, but when ion-ion repulsions are strong due to weak screening, transfer will be favored over uptake. Thus, as bulk concentrations increase, weak ion-ion repulsions decrease flux above the saturation point, while strong repulsions increase it.

Using a three-site model system, we untangle these two distinct profiles in response to increasing concentrations. When electrostatic interactions are excluded or screening is very strong, high ion occupancy can cause current to decrease above an optimal concentration. This is the consequence of release from the last binding site becoming rate limiting^21^ while uptake becomes more probable. In contrast, when ion-ion repulsion is strong and channel occupancy decreases, current asymptotically increases with increasing concentrations. While this behavior is more dramatic in the model system, it clearly mirrors the trends observed for flux between the ESP and NSP in the Shaker model. It is also consistent with experimental observations of decreasing flux above the ESP and previous explanations that it results from fast uptake relative to slow transfer.

This interplay between ionic repulsions and robust current under high concentrations could be a governing physical feature in the evolution of efficient ion channels. We also observe that variations in the dielectric constant influence the number of possible contributing pathways in ion channels, suggesting an alterative mechanism by which protein evolution could regulate electrochemical behavior.

Collectively, our findings emphasize the importance of retaining a full network description when studying the electrochemical behavior of ion channels. Simplification to a single or minimal set of pathways can lead to erroneous predictions for flux under different electrochemical conditions. Moreover, detailed insights into network behavior can optimize the use of exisiting experimental data and suggest new assays to discriminate between kinetic descriptions. Moving forward, the complementary strengths of MSMs and MsRKM offer promising opportunities. The former provides a wealth of information from molecular-level, bottom-up characterization while the latter provides a quantitative bridge to experimental data with complete network resolution of channel response to a wide range of electrochemical conditions. The integration of the two may drive significant advancements toward the development of kinetic modeling tools.

## AUTHOR INFORMATION

### Authors

**Hannah Weckel-Dahman** - Department of Chemistry, University of Utah, Salt Lake City, UT, 84112 – United States;

**Alex Daum** – Department of Chemistry, University of Utah, Salt Lake City, UT, 84112 – United States;

**Ryan Carlsen** – Department of Chemistry, University of Utah, Salt Lake City, UT, 84112 – United States;

**Tyler Southam** – Department of Chemistry, University of Utah, Salt Lake City, UT, 84112 – United States;

**Maxwell He** – Department of Computer Science, Northeastern University, Boston, MA, 02115 – United States;

### Notes

The authors declare no competing financial interest.

## ACKNOWLEDGMENT

This work was supported by NIH NIGMS (R35GM143117) and the computational resources provided by Expanse at the San Diego Supercomputing Center through the Advanced Cyberinfrastructure Coordination Ecosystem: Services and Support (ACCESS) program (allocation MCB200018) supported by NSF (grant nos. 2138259, 2138286, 2138307, 2137603, and 2138296), as well as the Center for High-Performance Computing (CHPC) at the University of Utah.

