## Supplemental Text for "Ion Channel Reaction Networks: Dielectric Screening and the Importance of Off-Pathway Flux"

### 1. SUPPLEMENTAL METHODS

#### 1.1. Optimization Schemes

*4S Model.* In the 4S Model wildtype optimization scheme, both global (Dual annealing) and local (Powell) optimization algorithms from the SciPy[1] python library were employed. All 56 rate coefficients were allowed to vary within their boundary conditions, with boundary conditions set as the diffusion limit ($1x10^{8}$ $mM^{-1}ms^{-1}$) for diffusion-limited binding events or with upper boundaries set at $1x10^{10}$ $ms^{-1}$ and lower boundaries set at $1x10^{-10}$ ($mM^{-1}$) $ms^{-1}$. Scipy's random number generator pulling from a log uniform distribution generated the initial seed rate coefficients for the wildtype optimization (See Figure S1A). The dual annealing algorithm was constrained to 25,000 non-consecutive optimization steps using a maxiter value of 25,000. Powell was constrained to a maximum number of function evaluations set to 100,000 (maxfev=100,000). All solutions did not reach the maximum number of function evaluations for the local function, with no more than ~92,000 steps observed before restarting the global algorithm. If an optimization run timed out before reaching 25,000 global searches or a solution, the run was restarted from the last optimization step completed. Unitary flux through the MsRKM model was compared to experimental values of the wildtype Shaker protein[2] using the residual function described below (See SI Section 1.2).

*4S-E Model.* The 4S-E model used the same Scipy parameters, algorithms, and steps as the 4S model, with the initial seed coefficients generated using Scipy's random number generator pulled from a log uniform distribution (Figure S1B). Boundary conditions for each base rate were identical to boundary conditions for the 4S model. The Δ values for each transition represent the $\Delta G^{\ddagger}_{elec}$ calculated according to Eq. 9, with optimization boundary conditions set using $\varepsilon'$=2 as the upper bound, representing the electrostatic repulsion expected for pure protein, and $\varepsilon'$=100 as the lower bound. To maintain the correct rate orderings, $\delta'_N$ multipliers were applied to Δ values for ion locations closer to the transition location. The lower bound for these multipliers was set to 1, guaranteeing that an ion moving closer to the transition location will not have less influence than an ion farther away. The upper bound was calculated as the ratio of two Δ values:

$$\delta'_N = \frac{\Delta_N}{\Delta_{N+1}} \tag{1}$$

For example, the upper bound for $\delta'_3 = \frac{\Delta_3}{\Delta_4} = \frac{(\frac{1}{r^T_{1,3}} - \frac{1}{r^R_{1,3}})}{(\frac{1}{r^T_{1,4}} - \frac{1}{r^R_{1,4}})}$.

Unitary flux through the MsRKM model was compared to the experimental values of the wildtype Shaker protein[2] using the same residual function as the 4S model.

*ESP and NSP Optimizations.* As the NSP appeared to be independent of the chosen dominant cycle, we selected four 4S-E solutions from each of the five dominant mechanistic pathways (20 total) for two additional optimizations. Both optimizations used the same protocol, detailed in SI 1.1, allowing only the off-pathway rate constants to change by up to +/- 10% of their original values to maintain the dominant pathway of each mechanism. In the first optimization, the rate constants were optimized to the same set of experimental conditions as the 4S-E model, but the original saturation IV curve (1.15 M) was fit to three new NSP concentrations, the first ranging from 506 mM to 9 M, the second the same NSP+5 M, and the third NSP+7 M. Above a base NSP of 9 M the optimization could no longer find a solution that met the acceptance criteria (<0.3 pA deviation). To test if the ESP was also dependent on the non-dominant pathways, an additional optimization procedure was performed to match the flux of the original ESP (506 mM) to a new ESP. The original experimental IV curves for 506 mM and 1.15 M of Heginbotham et al.[28] were mapped to new ESPs and new ESPs+0.5 M to enable a slight increase in flux between the two concentrations. The value of the ESP was then gradually changed between 350 mM and 3 M.

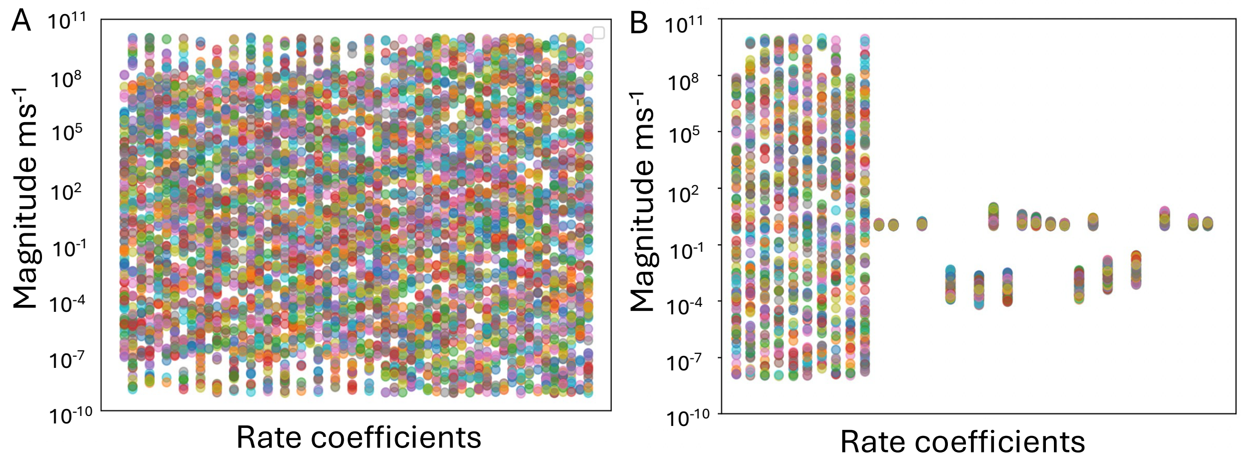


Figure S1. Seed rate coefficients randomly generated using a log uniform distribution for the A) 4S model and B) 4S-E model.

### 1.2 Additional Information

#### *1.2.1 System Considerations*

The Shaker channel is modeled with four $K^+$ binding sites (Figure 3) located at the S6 'gate' region ($S_1$)[3] and within the selectivity filter at T441($S_2$), G443($S_3$), and G445($S_4$)[4, 5]. The Shaker model assumes the open conformation of the protein and has no conformational constraints within the region of the selectivity filter.[6] Ions cannot tunnel (e.g., from S1 to S3) and the model assumes transitions are not coupled. This reduces the number of transitions from 240 (16*(16-1) possible transitions from 16 ($2^4$) states) to 56.

| **Model** | **Intra** | **TS 1** | **S 1** | **TS 2** | **S 2** | **TS 3** | **S 3** | **TS 4** | **Extra** |
|---|---|---|---|---|---|---|---|---|---|
| Base | -33 Å | -15 Å | -10 Å | -5 Å | 0 Å | 5 Å | 10 Å | 15 Å | 33 Å |
| All Shift | -33 Å | -17.5 Å | -15 Å | -10 Å | -5 Å | 0 Å | 5 Å | 12.5 Å | 33 Å |
| *Increment* | 0 Å | -.125 Å | -.25 Å | -.25 Å | -.25 Å | -.25 Å | -.25 Å | -.125 Å | 0 Å |

| | | | | | | | | | |
|---|---|---|---|---|---|---|---|---|---|
| Shift First Barrier | -33 Å | -17.5 Å | -10 Å | -5 Å | 0 Å | 5 Å | 10 Å | 15 Å | 33 Å |
| *Increment* | 0 Å | -.125 Å | 0 Å | 0 Å | 0 Å | 0 Å | 0 Å | 0 Å | 0 Å |
| Shift First Site | -33 Å | -17.5 Å | -15 Å | -5 Å | 0 Å | 5 Å | 10 Å | 15 Å | 33 Å |
| *Increment* | 0 Å | -.125 Å | -.25 Å | 0 Å | 0 Å | 0 Å | 0 Å | 0 Å | 0 Å |

Table S1 – Site locations for the 3S-E model. The base model was used for all models except those exploring the effect of site location on rectification. For models exploring the impact of site location on rectification, the maximum shift is shown in the first row, and the increments at which data was taken between the base location and the maximum shift shown are recorded in the increment row beneath.

*1.2.2 Experimental Data in the Residual Function*

Optimization runs minimize the loss function:

$$Objective = \frac{\Sigma_{ij}\left( \left(\mathcal{J}_{ij} - \mathcal{J}_{ji}\right) / \langle \mathcal{J}_{ij+ji} \rangle \right)^2}{\#_{ij}} + \Sigma_{sys} W\left(\mathcal{J}_{calc} - \mathcal{J}_{def}\right)^2 \quad (1)$$

where the first term captures microscopic reversibility in a mean squared error between forward and reverse flux ($\mathcal{J}_{ij}$ is the flux from state $i$ to state $j$) normalized over the total number of transitions ($\#_{ij}$) and the second term calculates the least squares difference between the calculated flow values ($\mathcal{J}_{calc}$) and the user-supplied values for every experiment ($\mathcal{J}_{def}$).

Only experimental data from single channel IV relationships in symmetrical solutions of $K^+$ were included in the loss function. For the WT optimization, six symmetrical $K^+$ concentration unitary I-V curves from L. Heginbotham et al. were used (43 mM, 73 mM, 206 mM, 325 mM, 605 mM, 1.15 mM), with the final concentration (1.15 M) used to predict saturation.[2] All experiments were weighted equally in the loss function.

#### 1.3 Voltage coupling profiles

The dimensionless coupling factor ($\varphi(z)$), also called the fraction of the membrane potential, serves to capture the normalized fraction of the voltage drop across a membrane that an ion would feel as it traverses from one side of the membrane to the other. In previous works, $\varphi(z)$ has been calculated using explicit solvent[7-10] or a linearized PB-V continuum approximation.[11] In this paper, all voltage coupling profiles used were constructed to represent a perfectly symmetrical coupling factor, as done in previous studies[12]. This approximation provides a useful description of the voltage profile along the z-axis of a simulated protein while removing any protein-specific voltage influence.

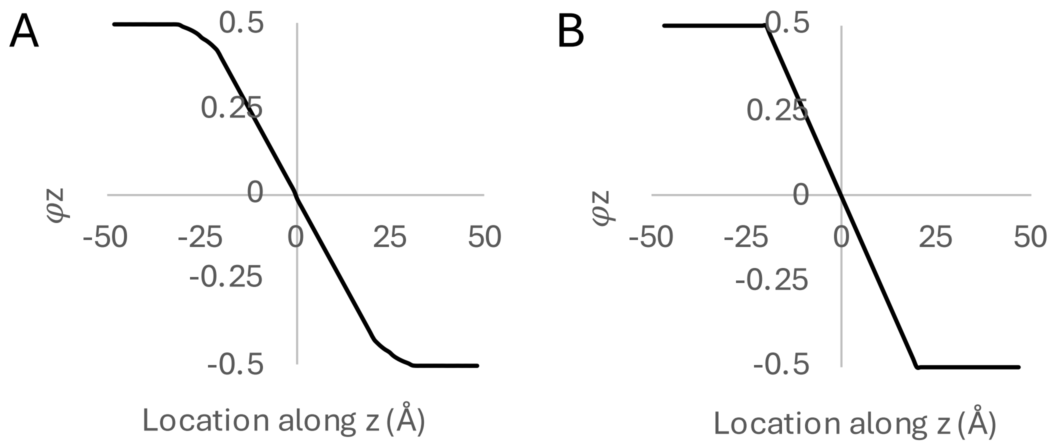


Figure S2. Voltage coupling profiles used in this study. A) Coupling profile designed to be perfectly symmetrical about 0, used with the 4S and 4S-E models. B) Coupling profile removing the taper between bulk and binding sites, used with the 3S-E model.

### 1.4 Cycle Visualization

Network graph visualizations were prepared in Gephi 0.10.0 with the Polygon Shaped Nodes plugin installed. A “Polygon” attribute with a value of 4 was given to each node to generate square nodes. Node size and edge weight values were calculated from the populations and flows, respectively, on a logarithmic scale to keep the smallest nodes and edges visible while still representing the larger nodes and edges in a manageable way.

Node sizes were calculated from the state populations using the following equation, with Gephi subsequently rescaling the node sizes to a minimum of 30 and a maximum of 60:

$$S = \log_{10}\left(\frac{p}{\min(P)}\right) + 0.5 \qquad \text{(S1.)}$$

where S is the node size, p is the respective state population, and P is the set of all populations in the network. Edge weight values were calculated using the following equation, with Gephi subsequently rescaling to a minimum weight of 2 and a maximum weight of 6, with an edge arrow size parameter of 4:

$$W = log_{10}\left(\frac{f}{\min\,(F)}\right) + 0.5 \qquad \text{(S2.)}$$

where W is the weight of the edge, f is the flow along the respective transition, and F is the set of all flows in the network. Individual node images were generated in BioRender. Node images and labels were edited into the network graph images and additional visual adjustments made using Inkscape.

### 2. SUPPLEMENTAL RESULTS

#### Supplemental Figures

| Transition | kij Base | Occupancy | kij Solution 2 | Electrostatic Forces | Vacuum kij |
|---|---|---|---|---|---|
| S1 binds 0000 -> 1000 | 1.06E+01 | S2 | 2.20E+04 | -1.01E-02 | 9.16E+00 |
| | | S3 | 7.73E+00 | -7.05E-03 | 9.57E+00 |
| | | S4 | 1.98E+04 | -5.34E-03 | 9.81E+00 |
| | | S2,S3 | 2.07E+03 | -1.71E-02 | 8.27E+00 |
| | | S3,S4 | 2.11E+05 | -1.24E-02 | 8.86E+00 |
| | | S2,S4 | 2.23E+05 | -1.54E-02 | 8.48E+00 |
| | | S2,S3,S4 | 1.78E+02 | -2.25E-02 | 7.65E+00 |

Table S2. Solution 3 versus coefficients derived using vacuum electrostatics. Implementation of electrostatics reveals incorrect rate relationships between previous solutions. Additional occupancies are expected to slow down the binding at the S1 site but Solution 2 optimized to faster transitions than possible.

| [100mM] ext: [100mM] int | 0.1V | 0.05V | 0V | -0.05V | -0.1V |
|---|---|---|---|---|---|
| Flux electrostatic model | -2.7566E+00 | -2.3100E-02 | 3.7048E-04 | 5.5760E-02 | 3.7235E+00 |
| Flux non-electrostatic model | -2.7566E+00 | -2.3100E-02 | 3.7048E-04 | 5.5760E-02 | 3.7235E+00 |

| 0V | [1mM]ext : [100mM]int | [25mM]ext : [75mM]int | [50mM] : [50mM] | [50mM]ext : [1mM]int | [100mM]ext : [1mM]int |
|---|---|---|---|---|---|
| Flux electrostatic model | -1.2370E-01 | -1.0170E-02 | 1.7522E-04 | 2.9100E-02 | 1.7017E-01 |
| Flux non-electrostatic model | -1.2370E-01 | -1.0170E-02 | 1.7522E-04 | 2.9100E-02 | 1.7017E-01 |

Table S3. Equivalency of the 4S and 4S-E Models

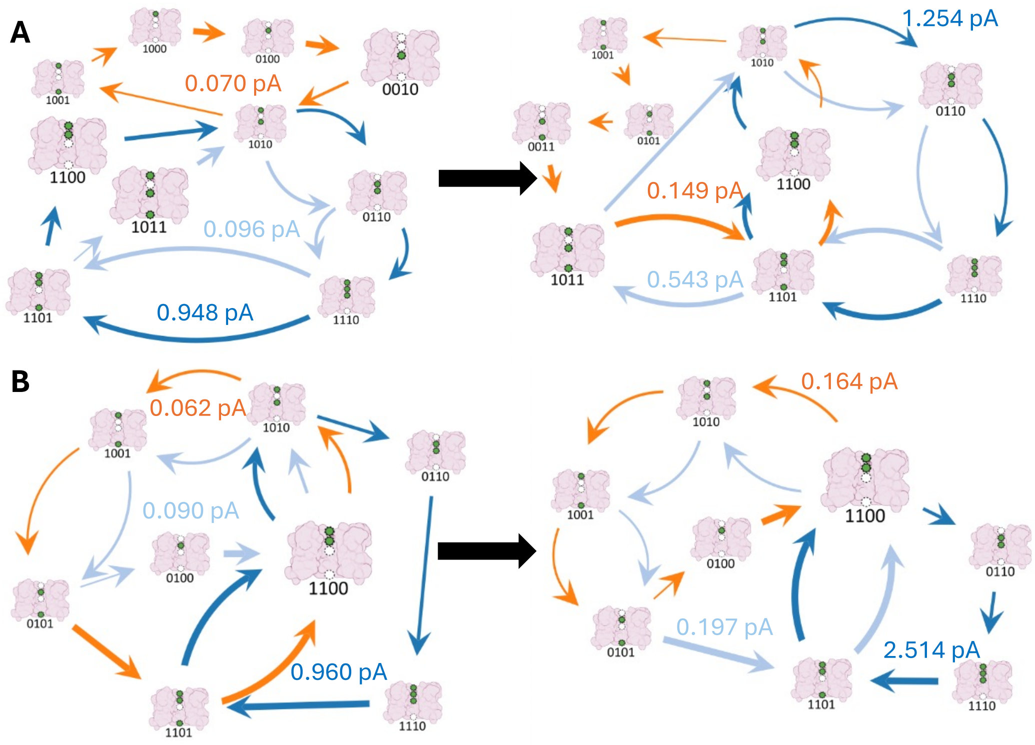


Figure S3. The top three flux-contributing pathways for Solution 9 (top) and Solution 12 (bottom) showing how the network changes from a 1 mM:50 mM concentration gradient (left) to a 1 mM:100 mM concentration gradient (right). Despite both solutions having the same dominant pathway, same RLS and same FLS, and similar contributions from the non-dominant pathways, we observe different network responses. SI Figure S3 was prepared using in-house graph visualization software based on the graph-tool python library.

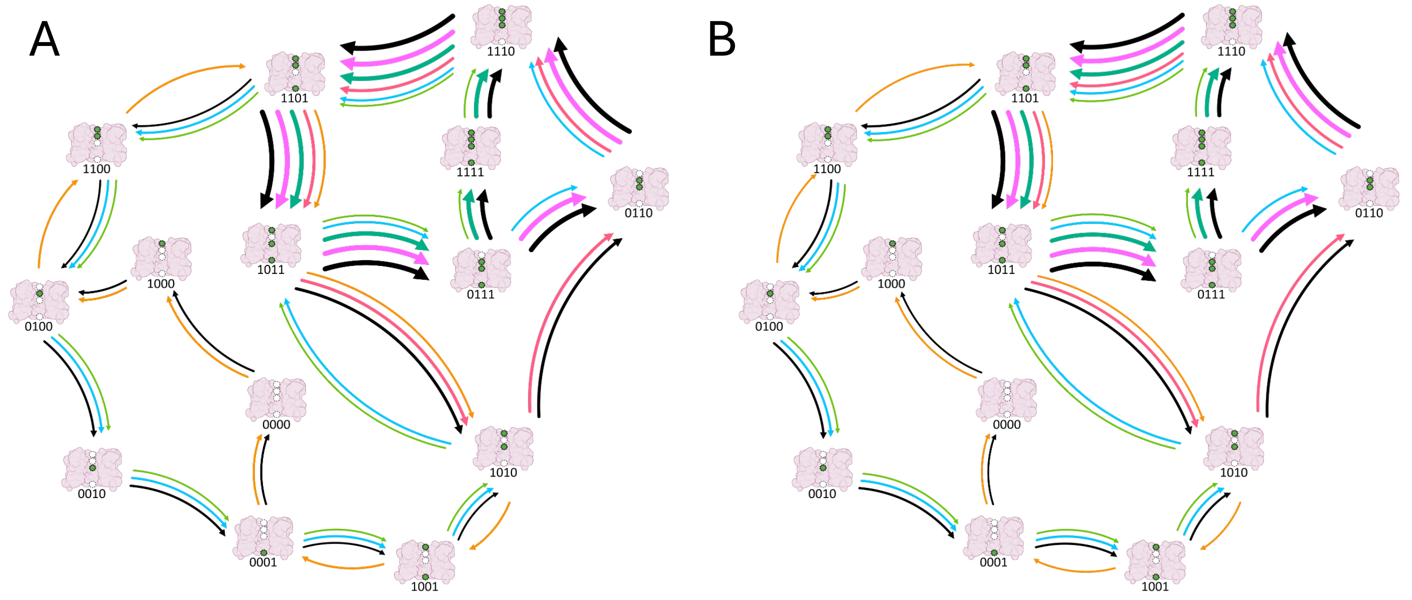


Figure S4. Network representation of WT Solution 78 at two concentration gradients greater than saturation conditions A) 5 M:1 mM [int]:[ext] and B) 7 M:1 mM [int]:[ext].

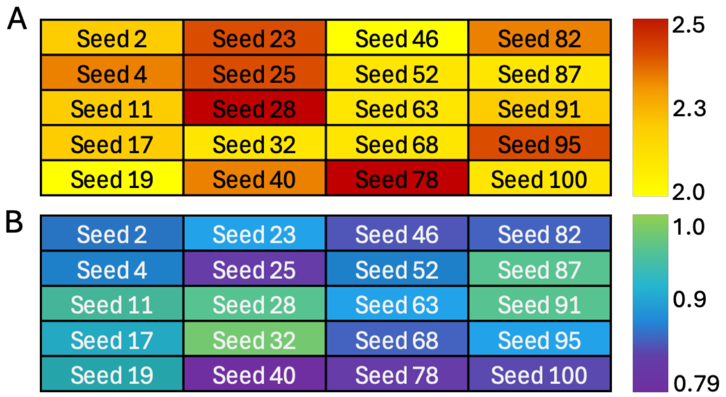


Figure S5. Tuning the saturation point by altering off-pathway cycles. A) Off-pathway cycles allow for a saturation point increase of 2.5x. B) Off-pathway cycles cannot as readily decrease the saturation point, with the maximum decrease observed at 0.79x.

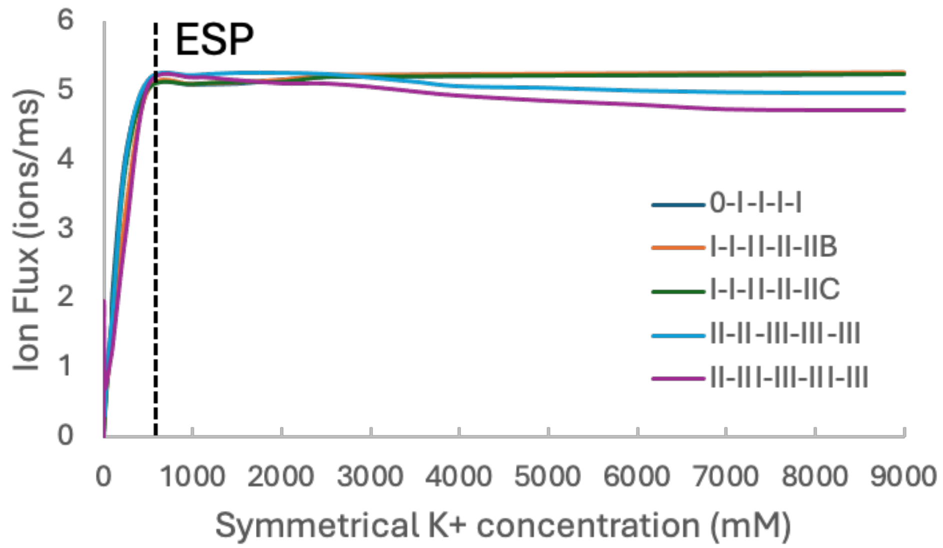


Figure S6. Difference in flux between the ESP (~506 mM) and the NSP for solutions with different dominant pathways. Pathways with one or two maximum ion occupancies in the channel (Dark blue, orange and Dark green) show gradually increasing net flux until the NSP (between 7 M and 8 M) was reached, while solutions with dominant pathways allowing a maximum of three ion occupancies in the channel show decreasing flux between the NSP and ESP.
